# The contribution of mitochondrial metagenomics to large-scale data mining and phylogenetic analysis of Coleoptera

**DOI:** 10.1101/280792

**Authors:** Benjamin Linard, Alex Crampton-Platt, Jérome Moriniere, Martijn J.T.N. Timmermans, Carmelo Andujar, Paula Arribas, Kirsten E. Miller, Julia Lipecki, Emeline Favreau, Amie Hunter, Carola Gómez-Rodríguez, Christopher Barton, Ruie Nie, Conrad P.D.T. Gillett, Thijmen Breeschoten, Ladislav Bocak, Alfried P. Vogler

## Abstract

A phylogenetic tree at the species level is still far off for highly diverse insect orders, including the Coleoptera, but the taxonomic breadth of public sequence databases is growing. In addition, new types of data may contribute to increasing taxon coverage, such as metagenomic shotgun sequencing for assembly of mitogenomes from bulk specimen samples. The current study explores the application of these techniques for large-scale efforts to build the tree of Coleoptera. We used shotgun data from 17 different ecological and taxonomic datasets (5 unpublished) to assemble a total of 1942 mitogenome contigs of >3000 bp. These sequences were combined into a single dataset together with all mitochondrial data available at GenBank, in addition to nuclear markers widely used in molecular phylogenetics. The resulting matrix of nearly 16000 species with two or more loci produced trees (RAxML) showing overall congruence with the Linnaean taxonomy at hierarchical levels from suborders to genera. We tested the role of full-length mitogenomes in stabilizing the tree from GenBank data, as mitogenomes might link terminals with non-overlapping gene representation. However, the mitogenome data were only partly useful in this respect, presumably because of the purely automated approach to assembly and gene delimitation, but improvements in future may be possible by using multiple assemblers and manual curation. In conclusion, the combination of data mining and metagenomic sequencing of bulk samples provided the largest phylogenetic tree of Coleoptera to date, which represents a summary of existing phylogenetic knowledge and a defensible tree of great utility, in particular for studies at the intra-familial level, despite some shortcomings for resolving basal nodes.

## Introduction

Current studies aiming at the construction of an ever more complete Tree-of-Life can draw on rapidly growing public taxonomic DNA databases (Chesters, 2017; Hinchliff et al., 2015; Hunt and Vogler, 2008; McMahon and Sanderson, 2006; Peters et al., 2011; Price and Bhattacharya, 2017; Zhou et al., 2016). Hitherto most taxonomic DNA data of higher eukaryotes have been obtained by sequencing from individual specimens, based on studies dedicated to particular target species. In insects, this approach requires focused collecting efforts and careful taxonomic identification, and may not achieve the broader goal of a fully sampled Tree. However, a rapid increase in taxon coverage is possible by direct sequencing of pooled multi-species DNA extracts obtained from mass-trapped arthropod specimens. Metagenomic sequencing of arthropods readily yields mitochondrial genome sequences that are assembled preferentially from the mixture of shotgun reads due to their high copy number (‘metagenome skimming’; Linard et al., 2016; Malé et al., 2014), allowing for species detection and phylogenetic placement of specimens in the samples. When focused on organelles, this approach is referred to as mitochondrial metagenomics or mito-metagenomics (MMG) (Crampton-Platt et al., 2016) and, if widely applied to samples from various habitats and biogeographic regions, may add large numbers of species to the assembly of the Tree-of-Life. The procedure is equally applicable to specimens that have not been formally described or characterized taxonomically, and thus even species that are new to science may be included in the tree, which can rapidly increase the taxonomic coverage especially in poorly known, species-rich and small-bodied groups of invertebrates.

In addition to greater species representation, mitochondrial genomes as those obtained through MMG are powerful markers to determine the phylogenetic position of the newly added taxa. Phylogenetic trees from MMG studies therefore may be more robustly supported than those resulting from conventional PCR-based analyses which usually only contain one or a few loci sequenced with ‘universal’ primers (Gómez-Rodríguez et al., 2015; Nie et al., 2017). In addition, existing studies have used different mitochondrial genes. Only the 658 bp barcode region of the *cox1* gene is now a widely agreed standard (Meusnier et al., 2008), while the representation of other genes in DNA databases varies greatly among taxa. Uneven gene coverage will result in data matrices with a high proportion of missing data and low phylogenetic support (Wiens, 1998). These matrices may be unable to define relationships among taxa represented by non-overlapping gene fragments (or provide poor support in case of partial overlap). However, the phylogenetic position of such taxa may still be approximated by establishing their relationships to a closely related mitogenome sequence, which determines their placement relative to each other indirectly through the mitochondrial loci shared with the full-length mitogenome. Thus, the fast production of mitogenomes from MMG could act as scaffolds for otherwise unmatched clades and complete the overall species representation and tree topology. Equally, these mitogenome sequences may represent species with already existing partial entries, which can be recognized for example by high sequence similarity to an existing *cox1* barcode. In these cases, the addition of MMG sequences can fill the missing data for that species and increase the phylogenetic signal of the matrix.

The present study attempts to integrate the growing amount of sequence data held at NCBI (GenBank) with newly generated mitogenome data from shotgun sequencing of specimen pools, to increase taxon coverage of the joint phylogenetic tree. We apply this approach to the arguably most diverse group of animals, the Coleoptera (beetles), whose total species richness is estimated to be in the millions (Basset et al., 2012), although currently the most comprehensive published DNA-based trees contain no more than 8000 species (Bocak et al., 2014). The Coleoptera has already been the focus of various MMG studies, e.g. to analyze basal relationship within the order and within several family-level taxa (e.g. Scarabaeidae, Chrysomelidae, and Curculionoidea) (Gillett et al., 2014; Gómez-Rodríguez et al., 2015; Nie et al., 2017), or to investigate species diversity in rainforests (Crampton-Platt et al., 2015), the soil (Andújar et al., 2015; Cicconardi et al., 2017), pollinators communities (Tang et al., 2015), and temperate terrestrial and aquatic habitats (Linard et al., 2016). This growing number of available mitogenomes provides a unique opportunity to draw together diverse samples for cross-phylogeny mitogenomic studies, and to link them to taxonomic data in GenBank.

To explore the contribution of the metagenomic approach for Tree-of-Life studies, we compiled public sequence data for the Coleoptera, and tested the effects of adding MMG data to the phylogenetic analysis. We assembled mitogenomes from several sets of shotgun sequence data from recent ecological and taxonomic studies, including those from poorly characterised tropical beetle assemblages obtained by mass trapping. A simple automated bioinformatics approach was used to integrated with data mined from the NCBI database. The analysis focuses on the contribution that MMG data make to the Tree and searched for their potential role as an anchor for linking non-overlapping mitochondrial gene sequences from public databases. The resulting tree of some 16,000 species constitutes the most complete compilation of phylogenetic data for the Coleoptera and represents a scaffold for forthcoming MMG and other data sources, which will iteratively enrich the tree of beetles.

## Material and Methods

### GenBank sequence extractions

Following the general scheme of the bioinformatics pipeline (Figure 1), selected loci were extracted from GenBank (Benson et al., 2013) using a set of steps previously developed by (Bocak et al., 2014). Briefly, a set of “bait” sequences of some 200 species covering the major lineages of Coleoptera was built for each target locus, whereby taxa were selected for completeness of available data and maximum clade coverage. Next, the Coleoptera fraction of the non-redundant (nr) database was retrieved to recover all potential homologous GenBank entries, by selecting all entries attached to the Coleoptera taxonomic identifier (taxid=7041). These sequences were converted to build a Blast searchable database, using the blastdb software. The above bait sequences were used as queries in Blastn (Camacho et al., 2009) searches (Evalue=1e-5, word size =12 and align_length>=60% of the expected locus length) against this database. The matching sequence regions were trimmed and extracted, building separate sets of sequences for each targeted locus. When several sequences were available for the same species (as identified in the NCBI taxonomy), only the longest fragment was kept. If several sequences of equal length were available, only the first-hit sequence was kept without consideration of subspecies or specimen information.

**Figure 1:**
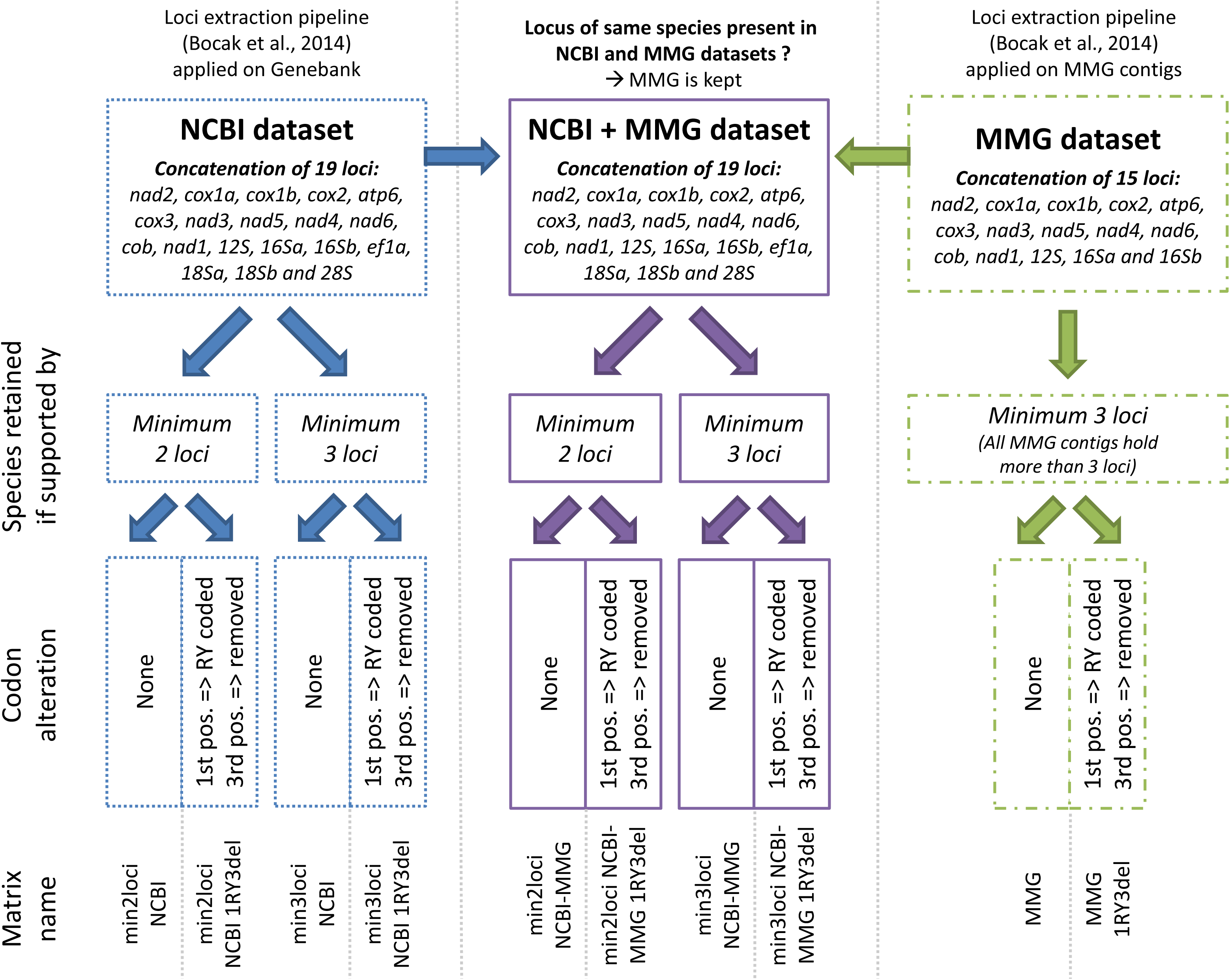
Sequence datasets built for this study and decision tree for generating the supermatices. Sets of loci were extracted from the GenBank and MMG contigs to respectively build the NCBI dataset (left, blue stippled lines) and the MMG dataset (right, green dashed lines). Both were merged in the MMG-NCBI dataset (center, purple solid lines). The loci support filters and sequence recoding applied to these datasets are reported and resulted in 10 different sequence matrices whose names are given at the bottom of the figure.

In total, three commonly used nuclear genes and 13 mitochondrial loci genes were retrieved (October 2015). The nuclear loci included a partial coding region of Elongation Factor 1-alpha (ef1a; ~600 bp), the 18S rRNA, split in two segments denoted 18Sa (~650 bp) and 18Sb (~1100 bp) and the 28S rRNA expansion segment (D2-D3 region, ~800 bp). All mitochondrial genes were included and the full length of each gene was used, except for *rrnL* (16S rRNA), for which two fragments were retrieved. These were denoted 16Sa and 16Sb, located between the V2-V3 (~300 bp) and V3-V4 regions (~220 bp) of the RNA gene. Additionally, the *cox1* gene was split into 2 segments (*cox1a, cox1b*) to accommodate the cox*1*-5’ 658 bp barcoding region and the widely represented *cox1-3’* portion bracketed by the Pat-Jerry primer pair (~700 bp). Thus, from these 16 genes we defined a total of 19 loci: *nad2, cox1a, cox1b, cox2, atp6, cox3, nad3, nad5, nad4, nad6, cob, nad1, rrnS, rrnLa, rrnLb*, ef1a, *18Sa, 18Sb and 28S.* The compiled data extracted in the above manner are referred to as ‘NCBI dataset’ (Fig. 1).

### Mitochondrial Metagenomics

Metagenomic shotgun reads were obtained from various existing studies motivated initially by separate phylogenetic, taxonomic, methodological or ecological questions (Table 1; see Suppl. File S1 for GenBank accessions). Out of these, the MT1 and MT2 datasets were partially based on long-range PCR of mitochondrial genomes or were Sanger sequenced NCBI data (273 mitogenomes of which 80 were by long-range-PCR (Timmermans et al., 2015b, 2010), whereas all other datasets were obtained by bulk shotgun sequencing of genomic DNA. Mitogenomes were extracted from these mixtures using the MMG protocol of (Crampton-Platt et al., 2016). The published datasets had been assembled and manually curated to various degrees previously, but under slightly different parameter settings for each set (different quality control, different assemblers). Here, raw data were used to re-build these contigs under uniform conditions and without any manual curation steps, to test the data quality achievable under a fully automated approach that is appropriate to large-scale application of MMG. A small proportion of these data had already been deposited at NCBI at the time of data extraction but they were removed from the NCBI mined dataset to avoid duplication. The shotgun mixtures underwent mitochondrial assembly starting from the corresponding raw Illumina libraries with the following protocol: (i) Libraries were trimmed to remove residual library adaptors with Trimmomatic v0.32 (Bolger et al., 2014) and low-quality reads were filtered with Prinseq v0.20.4 (Schmieder and Edwards, 2011). Only paired reads were retained. (ii) Mitochondrial reads were filtered with Blastn (Camacho et al., 2009) using the same reference database (i.e., all complete coleopteran mitochondria from GenBank, October 2015), to remove reads lacking similarity to mitogenomes. (iii) The filtered reads were then assembled with IDBA-UD v1.1.1 (Camacho et al., 2009; Peng et al., 2012) and only contigs of a minimum length of 3 kb were kept. (iv) Similarly to the GenBank sequence extraction, mitochondrial contigs underwent loci extraction through the custom pipeline (Bocak et al., 2014) for 15 mitochondrial loci (splitting *cox1a, cox1b* and *rrnLa and rrnLb*). The aligned matrix of contigs obtained in the above manner is referred to as ‘MMG dataset’ (Fig. 1). All parameters settings used in the assembly and data compilation are given in Suppl. File S1.

**Table 1:** Shotgun datasets used in this study and their origin.

| Dataset Identifier | Tree identifier | Publication | Initial study purpose | Sequencing approach | mitochondrial contigs | complete mitochondria (all coding genes) |
| --- | --- | --- | --- | --- | --- | --- |
| **B480** | B480 | Crampton-Platt et al., 2015 | Biodiversity | MMG | 155 | 85 |
| **BIUK** | BIUK | Crampton-Platt, PhD UCL 2015 | Biodiversity | MMG | 87 | 44 |
| **MT1** | JX* | Timmermans et al., 2010 | Phylogeny | Long-range PCR | 28 | 0 |
| **MT2** | JX* | Timmermans et al., 2016 | Phylogeny | Long-range PCR / MMG | 245 | 24 |
| **Curc** | Curc | Gillett et al., 2014 | Phylogeny | MMG | 110 | 74 |
| **Germany Barcode-of-Life** | amie | Present study | Phylogeny | MMG | 189 | 2 |
| **Chryso** | chryso | Gómez-Rodríguez, et al., 2015 | Ecology | MMG | 173 | 130 |
| **Staphylinidae** | staph | Present study | Phylogeny | MMG | 90 | 61 |
| **Soil** | soil | Andújar et al., 2015 | Ecology | MMG | 171 | 71 |
| **Ethanol** | 1VWB/2VGB | Linard et al., 2016 | Methodology | MMG | 45 | 29 |
| **M2015** | m2015 | Present study | Phylogeny | MMG | 33 | 21 |
| **Scarabaeidae** | scarab | Breeschoten et al., 2016 | Phylogeny | MMG | 86 | 17 |
| **ChrysoT** | chrysot | Rui et al., submitted | Phylogeny | MMG | 72 | 49 |
| **French Guyana** | FG | Present study | Biodiversity | MMG | 153 | 71 |
| **Panama** | panama | Present study | Biodiversity | MMG | 236 | 100 |
| **Scolytinae** | scoly | Milller et al., 2015 | Methodology | MMG | 63 | 53 |
| **Cocci** | cocci | Paula et al., 2014 | Ecology | MMG | 6 | 6 |

### Multiple alignments and supermatrix

Eight mitochondrial genomes from the supraorder Neuropterida, the presumed sister group of Coleoptera (+ Strepsiptera), were added as an outgroup to each locus (Suppl. File S1). All protein-coding loci from the NCBI and MMG datasets were aligned with transalign (Bininda-Emonds, 2005; Sievers et al., 2011), except for the larger *cox1a* and *cox1b* datasets (more than 40,000 sequences each), which were aligned with Clustal Omega (Sievers et al., 2011) that is more suitable to the alignment of large numbers of sequences, using the ‘-auto’ configuration and default parameters. Each alignment was then manually curated and trimmed to start with the first and end with the last base of a codon. During this editing step, all gene boundaries extending beyond the average ORF length were removed. All rRNA fragments (*rrnS, rrnLa, rrnLb, 18Sa, 18Sb, 28S*) were aligned in two steps. A first multiple alignment was built with Clustal Omega using default parameters, then realigned with Mafft v7.123b (Katoh and Standley, 2013) using the FFT-NS-2 method, a gap extension value of 1.0 (option -gep) to favor longer indels in the conserved regions, and other parameters set to default.

All alignments were grouped under three concatenation schemes (Figure 1): (i) ‘NCBI’ based on the 19 concatenated loci (15 mitochondrial + 4 nuclear) of the ‘NCBI dataset’; (ii) ‘MMG’ containing the 15 concatenated mitochondrial loci from the ‘MMG dataset’ (also including the Timmermans et al., 2015b) data; see above); and (iii) ‘NCBI+MMG’ containing 15 mitochondrial + 4 nuclear concatenated loci, merging the two original datasets (Fig. 1, center). For the latter, if NCBI and MMG loci were both available for the same species (identified by an exact match of an NCBI entry to any portion of the mitogenome), only the MMG-based sequence was kept. For the NCBI-only and NCBI+MMG datasets we built 4 separate matrices (Fig. 1) requiring either a minimum of 2 or 3 available loci for each taxon (denoted min2loci and min3loci), and either using all nucleotides or removing the 3^rd^ codon positions and RY coding of 1^st^ positions (denoted 1RY/3del). All mitogenomes included more than 2 loci, and thus for the MMG-only dataset we conducted searches on all available contigs. Note that genetic variants in pooled DNA extracts generally collapse into a single contig during assembly (Gómez-Rodríguez et al., 2016) and therefore separate contigs were considered equivalent to different species.

### Tree labelling and taxonomic retention index

Leaves on the tree were labelled extensively (see Suppl. File S1 for details) with (i) the locus code, representing which of the 19 loci are available for a particular species in the matrix, (ii) the taxonomic code, a shorthand alphanumeric code that compresses the Linnaean taxonomy associated to the species (see Suppl. FIle S2 for complete list), (iii) the full Linnaean binomial as given on GenBank or from other sources, at the lowest taxonomic level available, and (iv) only for terminals based on MMG contigs, a short metadata string indicating the source dataset (Table 1), a contig identifier, the keyword “MERGED” when concatenated with GenBank data, in particular for nuclear genes, followed by a keyword referring to the method used to attribute a species identification to the MMG contig.

The locus code (see Suppl. File S1) is a vector of 19 positions either marked by ‘X’ (locus present in the supermatrix) or ‘-’ (locus absent). The position of each symbol was given in the canonical mitochondrial gene order: *nad2, cox1a, cox1b, cox2, atp6, cox3, nad3, nad5, nad4, nad6, cob, nad1, rrnLa, rrnLb, rrnS*, followed by the 4 nuclear loci in the order: ef1a, 18Sa, 18Sb, 28S. For instance, a common locus string is −XX X, indicating a species supported by a complete *cox1* gene (loci *cox1a*+*cox1b*, in positions 2 and 3) and a 16S RNA fragment (*rrnLa*, position 13). Similarly, the code --XXXXXXXXXXXXX—X indicates a species supported by a nearly complete mitochondrial contig (only *nad2* and *cox1a* are missing) complemented by a 28S RNA sequence extracted from NCBI.

The taxonomic code compresses the Linnaean taxonomy associated to the sequence according to the GenBank classification (Federhen, 2012) for seven hierarchical levels: suborder, infraorder, superfamily, family, subfamily, tribe, genus, species. Names at each level were abbreviated to include only the first or second letter, with a third letter added to discriminate among names with the same starting letters. After each abbreviated name the hierarchical levels were given (1 to 7 from highest to lowest taxonomic level). The alphanumeric codes were also used as pseudocharacters to calculate the taxonomic Retention Index (tRI) at each hierarchical level, as a measure of the fit of the tree to the Linnaean classification (Hunt and Vogler, 2008).

The identification labels of assembled shotgun mitogenomes were based on identifications with the *cox1* barcodes generated during the original MMG studies (Table 1). Briefly, internal barcodes were produced for specimens prior to DNA mixing for bulk shotgun sequencing and were used as ‘bait sequences’ to link specimen and mitogenome (see Crampton-Platt et al., 2016). When barcodes were not available, contigs were identified to species level by Blast matches (>99% identity) to (i) *cox1* records from NCBI Genbank and (ii) the BOLD database, or (iii) where no such identification via existing entries was available, contigs were labelled at family level, based on their phylogenetic position in preliminary tree searches. The taxon label includes the method of identification (annotated from barcodes or from tree topology).

### Tree searches

Phylogenetic trees were generated using likelihood searches under data partitioning. Mitochondrial genes were attributed to a total of 7 data partitions, one for each codon position of protein coding genes on the forward (3 partitions) and reverse (3 partitions) strands, plus one partition for the two rRNA genes. Each nuclear gene was assigned to a single additional partition (ef1a, 18Sa+18Sb, 28S), for a total of 10 partitions. Using both the CIPRES web server (Miller et al., 2010) and in-house servers, tree searches were performed with RAxML-HPC2 v8.2.6 (Stamatakis, 2014) under the GTRCAT model of nucleotide substitution, which approximates a GTR+Γ model at reduced computational cost, and the likelihoods of the final tree topologies were re-evaluated under GTR+Γ. For the different tree searches (NCBI-only, NCBI+MMG, MMG-only) based on a minimum of three loci (min3loci, see Suppl. File S1), 20 independent runs with different starting seeds were performed, and the tree with the best likelihood score was selected. For searches on the larger datasets with a minimum of 2 loci (min2loci), we used the min3loci tree as starting tree (RAxML option --t). This starting tree option did not constrain the tree search, but started the taxon addition from a given topology of the smaller dataset. In addition, tree searches were performed under RY coding of 1^st^ and removal of 3^rd^ codon positions (1-RY/3-del), to reduce long-branch attraction due to compositional and rate heterogeneity. We did not obtain measures of node support (e.g. bootstrap values) due to the computing limitations imposed by the size of the studied matrices.

## Results

### NCBI and MMG extractions

The NCBI extraction revealed a preponderance of *cox1* sequences representing more than 46,000 species for each segment (*cox1a, cox1b*). The next highest numbers were for the *rrnL* segments (*16S*a and *16S*b) with over 13,000 species, and the *cox2, cytb* and 18S rRNA genes, available for around 5,000 species each. These loci represented a great diversity at all taxonomic levels (Figure 2A, left). Other mitochondrial and nuclear loci were underrepresented, restricted to a few clades, and on average corresponded to sequence fragments shorter than the full-length coleopteran gene, which was particularly striking for the widely used *cox3, nad5* and *cytb* genes (Figure 2A, bottom-right histogram). Changes in the stringency of the Blast searches using e-values of e=-12 to e=+1 showed little differences in the number of sequences obtained, except under the least stringent conditions (Suppl. Table S1).

**Figure 2:**
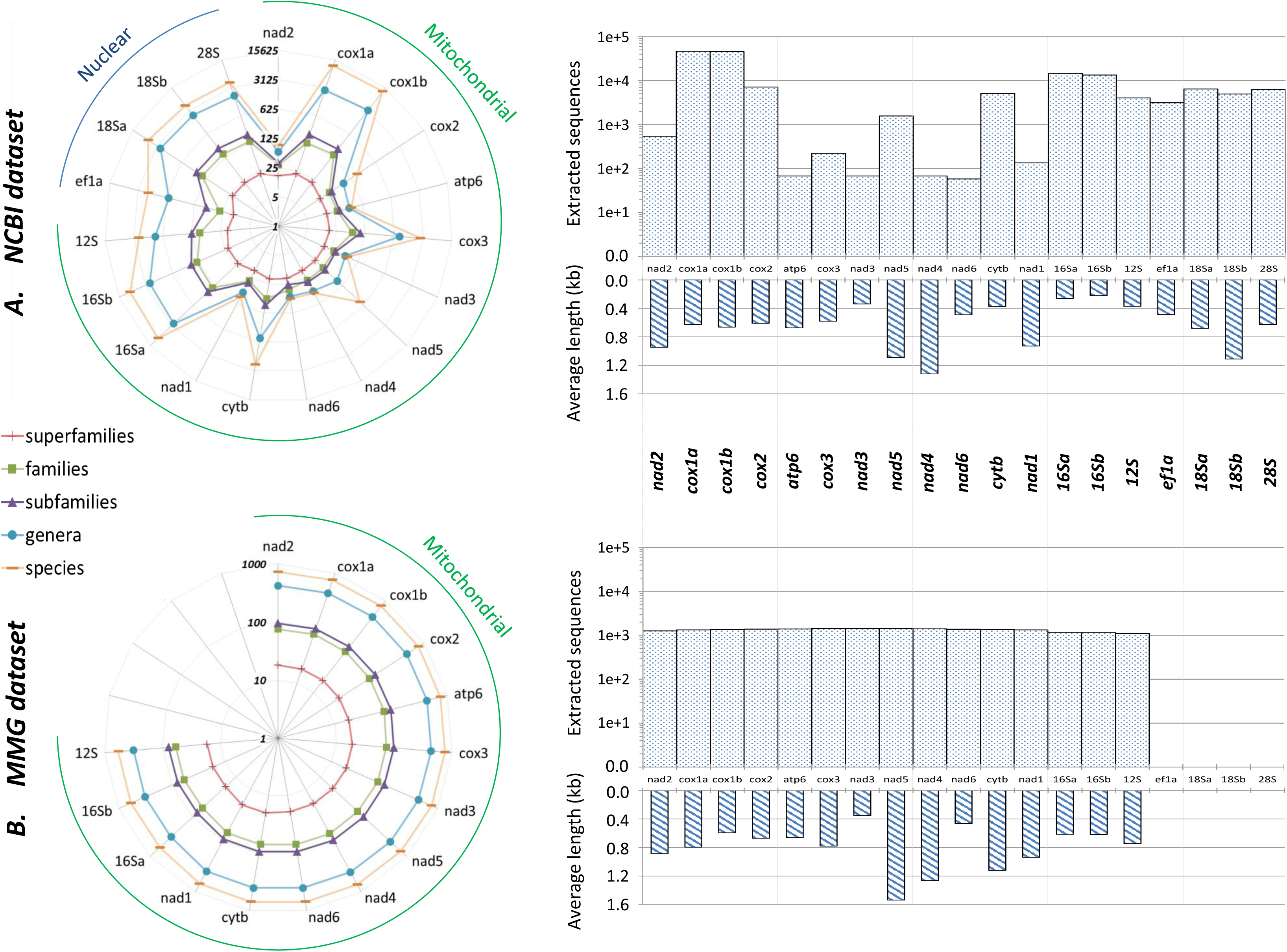
Composition of the extracted sequence data. Loci statistics are reported for (A.) the NCBI extraction (split in 15 mitochondrial and 4 nuclear loci) and (B.) the MMG datasets (15 mitochondrial loci). Left radars illustrate the taxonomic evenness of the extraction, with one unit corresponding to one clade of the following taxonomic levels: species (orange dashes), genera (blue dots), subfamilies (purple triangles), families (green squares) and superfamilies (red crosses). On the right side, dotted histograms report the total number of entries retrieved for each locus (including multiple entries belonging to the same species) on a log scale, while dashed histograms reports the corresponding average sequence length.

The mitogenome assembly conducted independently on each of the 17 shotgun datasets produced between 6 and 236 contigs, for a total of 1942 mitochondrial contigs, of which 561 contigs were obtained from previously unpublished datasets (Table 1). The contigs included complete (as indicated by their >15kb length) and partial mitochondrial genomes. For most shotgun datasets, a complete set of protein coding and rRNA genes was obtained in one half to two thirds of contigs, but with the notable exception of one dataset (Table 1). The distribution of recovered mitochondrial loci (Figure 2B, upper-right histogram) and their representation at all hierarchical levels were largely uniform (Figure 2B, left). The MMG-derived gene extractions were mostly composed of the expected full length sequences, in contrast to the partial genes in the NCBI dataset (Figure 2B, lower-right histogram). After assigning taxonomic names (see Materials and Methods), MMG contigs were complemented with NCBI data for the same species, adding the corresponding nuclear loci (and mitochondrial loci missing in incomplete contigs). In the min2loci NCBI+MMG matrix a total of 666 terminals were a combination of both sources, i.e. for ~1/3 of all MMG mitogenomes (see Suppl. File S1 for a list of all combined terminals). In many cases only the *cox1* sequences were available at NCBI, and were consequently discarded in favor of the longer MMG contig (see Materials and Methods).

### Taxonomic composition

The NCBI extraction recovered 165 out of 189 currently recognized beetle families, including 257 subfamilies, 5441 genera and 30,700 species (Table 2). After concatenation, 13,666 species were omitted from the tree searches on the final dataset because they did not have sequences for a minimum of 2 or 3 of the target loci and were mainly represented by *cox1* and *rrnL* fragments (data not shown). The minimum locus requirement mainly removed species at lower hierarchical levels, e.g. retaining only 39.0% and 52.1% of the total species available at NCBI for the min3loci and min2loci datasets, respectively, while retaining 95.8% and 97.6% of family level taxa (corresponding to 7 and 4 unrepresented families). The MMG set contained 120 families and 139 subfamilies, which is a minimum estimate, as many sequences remained unidentified. Both datasets overlapped in the coverage of higher taxa present but the proportion of taxa in the MMG set also represented in the NCBI extraction decreased towards the tip (species) level from 54.5% at the family level to a mere 3.4% at the species level (Table 2). At the same time, the MMG data contributed an increasing number of unique taxa for each hierarchical level. After redundancy correction for the species represented by both datasets, the total species representation in the min3loci and min2loci datasets was 11944 and 15983 terminals, respectively, including 1942 MMG contigs (Table 2).

**Table 2:**
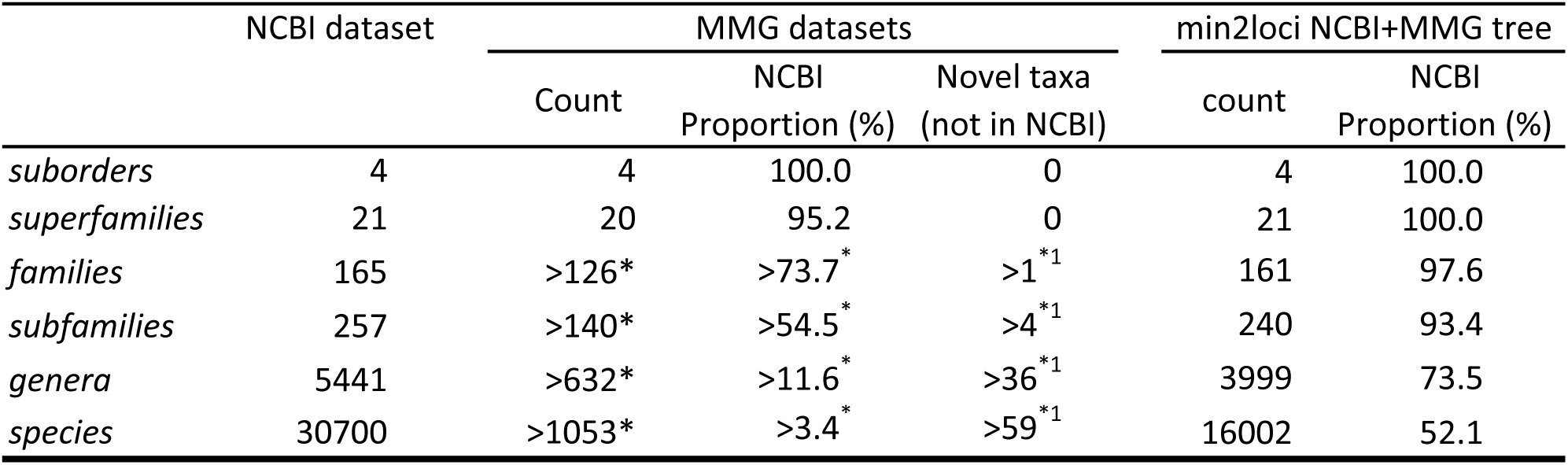
Taxonomic content of the NCBI, MMG and NCBI+MMG coleopteran supermatrices. “NCBI Proportion” describes the proportion of the NCBI coleopteran taxonomy covered by each dataset. * Numbers are low estimates as many contigs belongs to currently unidentified coleopteran species. ^1^ Mitochondrial contigs belonging to clades which were not barcoded in BOLD or defined in the NCBI taxonomy in October 2015 (date of the NCBI data extraction).

### Phylogenetic analyses

Tree searches were conducted on five datasets; the MMG contigs, the NCBI-only database requiring a minimum of either 3 or 2 loci, and the combined MMG and NCBI database with minimum of 3 or 2 loci (the min3loci and min2loci databases). Except for MMG-only, these datasets included >10,000 terminals with a large proportion of missing data in many sites. We first assessed the effect of different starting trees on the likelihood of the trees obtained with the RAxML software, launching 20 tree searches under different starting seeds on the min3loci dataset with all nucleotides partitioned into 10 partitions (Material and Methods). Each search took 5 to 9 days of calculation using 12 or 16 CPU cores on the Cipres web server and in-house server launches, respectively. The best tree had a likelihood score of −11,449,869.61, with other trees worse by between −263.00 and −1046.15 log likelihood units (final likelihoods were calculated under GTR+Γ on each topology for direct comparisons). The resulting tree topologies were overall similar although they differed in various places, including the basal relationships of the polyphagan superfamilies (see Suppl. File S1 for an example).

Tree topologies were assessed with the tRI, a measure of fit to the Linnaean taxonomy, at four hierarchical levels (genus, subfamily, family, superfamily) (Table 3). The NCBI min3loci (10,063 taxa) and MMG+NCBI min3loci (pruned, 10,063 taxa) datasets showed the highest values at all four levels (Table 3, bold values), in particular for superfamilies, whose distributions were almost entirely consistent with the tree (tRI=0.996 and 0.997). The 1-RY/3-del coding (Table 3: trees T1, T3, T5, T7) produced the exact same or very similar tRI values at high taxonomic levels, but these dropped slightly at the subfamily and particularly genus level. The tRI values for MMG were much lower at all hierarchical levels, ranging between 0.650 and 0.693, except for superfamilies (tRI=0.963) (Table 3: T8, T9). These values were directly correlated to the completeness of taxonomical annotation of the different studies from which these datasets were obtained. For example, several ecological MMG datasets (Panama, French Guyana, Soil) had been fully identified at superfamily level but not necessarily at lower taxonomic levels. When NCBI and MMG data were analyzed together, whether using the min3loci or min2loci datasets, the values were slightly reduced compared to the tRIs from the NCBI data alone (Table 3: T4 and T5). However, after pruning of the MMG data from the tree, the original values were largely regained or even slightly increased (Table 3: T4 MMG-pruned and T5 MMG-pruned), indicating a similar phenomenon of signal dilution.

**Table 3:** Taxonomic retention index of the NCBI, MMG and NCBI+MMG coleopteran trees. The taxonomic retention index (tRI) was calculated for four taxonomic level: genus, subfamily, family and superfamily. The 2 trees with best tRI scores are highlighted in bold. Asterisks mark MMG+NCBI trees resulting to less terminals than the summed of NCBI and MMG terminals, because of merged terminals (same species).

| Tree name | NCBI trees |  |  |  | NCBI+MMG trees |  |  |  |  |  |  |  | MMG trees |  |
| --- | --- | --- | --- | --- | --- | --- | --- | --- | --- | --- | --- | --- | --- | --- |
|  | min3loci |  | min2loci |  | min3loci |  |  |  | min2loci |  |  |  |  |  |
|  | T0 | T1 | T2 | T3 | T4 | T4, MMG<br>pruned | T5 | T5, MMG<br>pruned | T6 | T6, MMG<br>pruned | T7 | T7, MMG<br>pruned | T8 | T9 |
| **Codon recoding** | none | 1RY3del | none | 1RY3d | none | none | 1RY3d | 1RY3d | none | none | 1RY3d | 1RY3d | none | 1RY3d |
| **terminals** | 10063 | 10062 | 14612 | 14612 | 11944 | 10063 | 11942 | 10062 | 15983 | 14028* | 15983 | 14028* | 1876 | 1876 |
| **tRI genus** | **0.868** | 0.854 | 0.819 | 0.785 | 0.820 | **0.867** | 0.803 | **0.851** | 0.782 | 0.824 | 0.754 | 0.798 | 0.693 | 0.693 |
| **tRI subfamily** | **0.946** | 0.941 | 0.930 | 0.922 | 0.857 | **0.950** | 0.855 | **0.947** | 0.860 | 0.933 | 0.856 | 0.929 | 0.656 | 0.653 |
| **tRI family** | **0.986** | 0.985 | 0.977 | 0.977 | 0.896 | **0.987** | 0.896 | **0.986** | 0.910 | 0.982 | 0.910 | 0.980 | 0.650 | 0.663 |
| **tRI superfamily** | **0.995** | 0.995 | 0.991 | 0.991 | 0.942 | **0.996** | 0.942 | **0.997** | 0.953 | 0.995 | 0.953 | 0.995 | 0.953 | 0.963 |

The largest MMG+NCBI tree (T4) was inspected for the details of the topology, as the most complete summary of coleopteran relationships obtainable from the current NCBI database (Fig. 3; Suppl. File S4). We made extensive comments on the tree topology in the ‘small’ suborders Myxophaga, Archostemata and Adephaga with 25, 9 and 3,406 terminals, respectively, in the light of existing taxonomic knowledge and previous studies, including those that generated these data (Suppl. File S1). The subclades of the tree in some cases were almost entirely from entries of a single phylogenetic study (e.g. in Gyrinoidea), whereas other subclades combined the results from multiple studies and thus extended the taxon sampling in this tree beyond any existing work (e.g. in Cicindelidae). We found a general agreement with the expected relationships, while inconsistencies mostly affected individual terminals or small subgroups that were apparently misplaced (see Suppl. File S1 for detailed discussion). Likewise, the expected major clades and basal relationships of Polyphaga, including almost the entire set of infraorders and superfamilies (Mckenna et al., 2015; Timmermans et al., 2015b; Zhang et al., 2018), were recovered (Fig. 3; Suppl. File S4). Basal relationships within Polyphaga showed the basal split of Scirtoidea from the ‘core Polyphaga’ (Hunt et al., 2007; Zhang et al., 2018), and the branching of the series (infraorder) Elateriformia (Byrrhoidea, Elateroidea), followed by the Staphyliniformia plus Scarabaeiformia (Staphylinoidea + Hydrophiloidea + Scarabaeoidea), and the Bostrichiformia as sister to the large series Cucujiformia, which are arranged in a plausible topology (((Tenebrionoidea (Cleroidea (Cucujoidea (Chrysomeloidea, Curculionoidea). However, closer inspection of the tree (Fig. S4) also revealed individual species or smaller sublineages to be removed from the main clades representing a higher taxon, which was reflected in the tRI of <1.0. This is particularly evident in the two small suborders which were not monophyletic with respect to each other, presumably due to attraction of taxa with overlapping gene coverage, while the difficult basal relationships of the four suborders also were not in agreement with transcriptome data (Misof et al., 2014). Comparing the topologies from the min2loci NCBI-only and NCBI+MMG datasets (under 1-RY/3-del) many of the major lineages were monophyletic in both trees, but two groups (Bostrichiforma and Staphylinoidea) were recovered only with the MMG data included.

**Figure 3:**
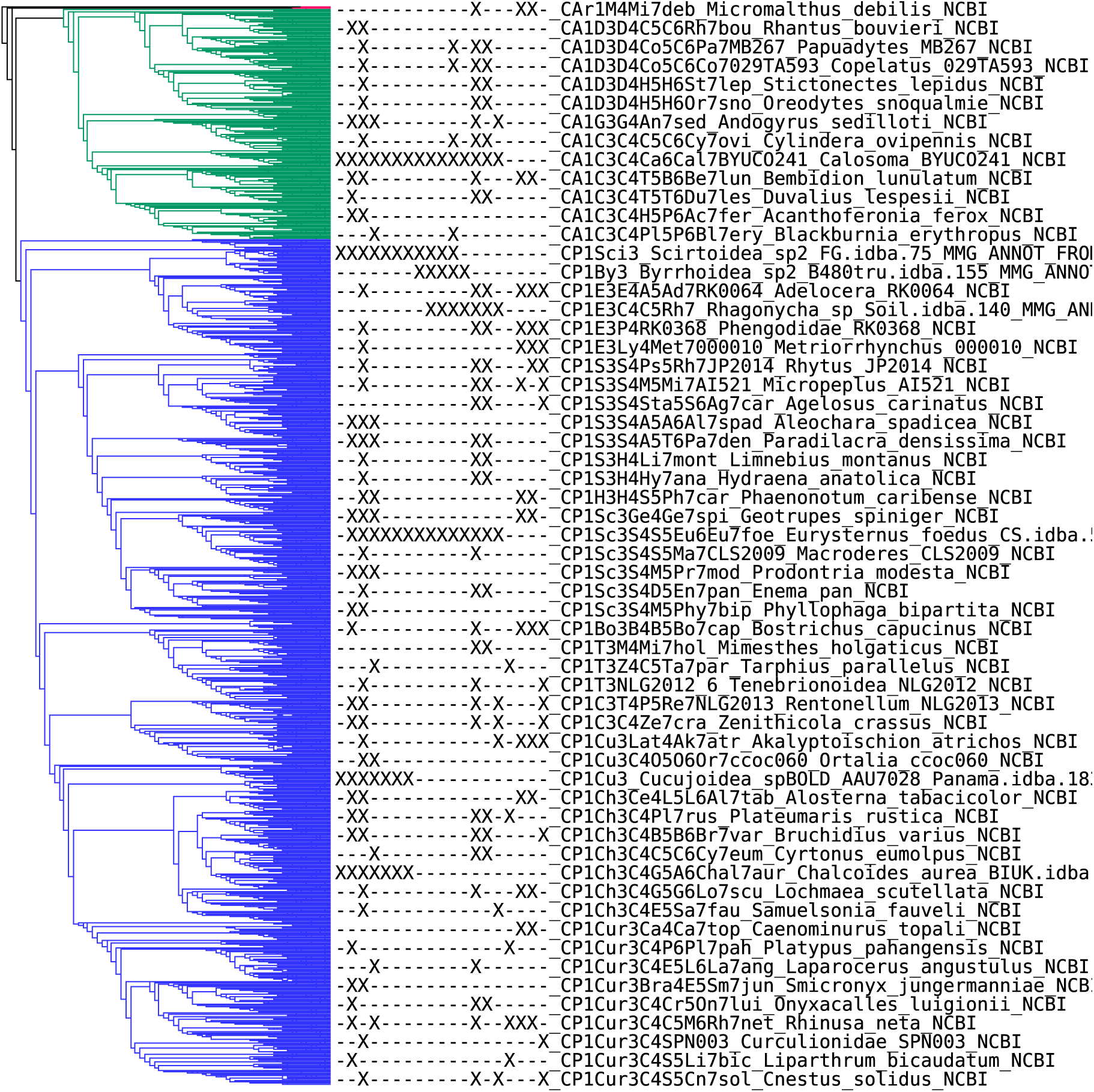
A graphical representation of the NCBI+MMG tree. This is a condensed version of the full tree of 15983 terminals requiring a minimum of two loci under 1RY/3del coding. The taxon labels were created with the Dendroscope tree viewer that displays sparse labels upon condensing of the tree. The complete unfolded tree is presented in Suppl. File S4. The naming scheme for terminals is according to the locus and name codes described in Material and Methods and Suppl. File S1. Ar1 Archostemata (red); A1, Adephaga (green); P1, Polyphaga (blue); C3, Caraboidea; D3, Dytiscoidea; Sci3, Scirtoidea; E3, Elateroidea; B3, Byrrhoidea; S3, Staphylinoidea; H3, Hydrophiloidea; Sc3, Scarabaeoidea; Bo, Bostrichoidea; T3, Tenebrionoidea; C3, Cleroidea; Cu3, Cucujoidea; Ch3, Chrysomeloidea; Cur3, Curculionoidea. See Suppl. File S2 for detailed classification key.

## Discussion

### MMG approaches and the Coleoptera Tree-of-Life

Phylogenetically informed studies of species diversity rely increasingly on genomic approaches (Maddison, 2016). While species-level trees already exist for major vertebrate phyla, and ambitious plans for full-genome sequencing of all species have been widely articulated (Zhang et al., 2014), the species-level phylogenetic analysis of a group of insects containing potentially millions of species has not been seriously considered. The MMG technology has the potential to provide rapid species recognition and phylogenetic placement of numerous species at the same time, thus setting the context for studies of evolutionary biology and ecology of the most diverse groups of animals (Crampton-Platt et al. 2016). With >1,900 contigs (representing nearly this number of different species, including 1000 fully identified species in 630 genera) the current database greatly exceeds the ~250 beetle mitogenomes previously available. In addition, this type of data will rapidly expand in the near future, not least because of the possibility of hybrid enrichment of mitochondrial genomes (Liu et al., 2016; Zhang et al., 2014) and improved sequencing methods that ensure efficient assembly (Mostovoy et al., 2016).

The inclusion of numerous mitogenome sequences into a highly incomplete matrix might improve the tree topologies and increase nodal support by reducing the overall proportion of missing data. They might also act as a scaffold to non-overlapping mitochondrial fragments from the NCBI entries. However, the improvements of tree topologies after inclusion of the MMG mitogenomes were negligible according to the test applied here. The comparison is based on the tRI values, i.e. a measure that integrates taxonomic information in relation to the Linnaean classification over the entire tree. The Linnaean taxonomy seems to be an excellent fit to the tree obtained from the NCBI data, in particular at the highest levels (suborder and superfamily), indicating that the tRI is a good measure of tree quality. Yet, the MMG data did not perform well by this criterion. The interpretation of this result should consider that the mitogenome data were obtained in the course of studies of various lineages and ecosystems unrelated to the aims of the current work. Three of the largest datasets (Soil, Panama and FG) represent specimens identified based on close matches to DNA barcodes and GenBank entries, or based on in-house sequences from specimens identified with various degrees of confidence. Many of these identifications are incompletely defined, e. g. only at the superfamily level, which would reduce the per-character tRI due to lack of data, as consistency with the tree may not be recognizable. High levels of ‘missing data’ in the calculation of the tRI can reduce the measure of consistency with the tree (Lanfear et al., 2014; Maddison, 1993). In addition, some names may be incorrect, creating conflict with the taxonomy. Therefore the low tRI from the MMG data alone (Table 3) is not a meaningful representation of the phylogenetic power of these mitogenomes. More relevant is the fact that the NCBI+MMG combined tree shows high tRI, and after pruning the MMG data from this combined tree (to avoid the annotation effect described above) the pruned NCBI+MMG tree even shows a slightly higher tRI, i.e. the MMG data improved the tree topology obtained from the NCBI data.

The MMG data might be further strengthened with more uniform taxon selection, which differed greatly across various parts of the tree. For example, among 442 species of Cicindelidae (tiger beetles) only one species is represented by a mitogenome, whereas 1,649 species of Chrysomelidae (leaf beetles) include nearly 300 mitogenomes. If applied more widely, MMG can could easily contribute more uniform coverage and might stabilize the tree, as hypothesized. Second, the MMG data were obtained with fully automated procedures for mitogenome assembly and matrix construction, as the approach is designed for high-throughput analysis of biodiversity samples. Several steps of data curation including the use of multiple assemblers and super-assembly of the products generally result in longer contigs and help to recognize chimeras and other spurious portions mostly near the ends of a contig (Gomez-Rodriguez et al., 2015). These incorrect terminal portions may also be responsible for the split of partial contigs that would otherwise be fused, which are evident in the tree as closely related contigs with marginally overlapping locus strings (Suppl. File S1). In addition, the quality of the MMG data is further affected by the gene extractions performed using Blast (in analogy to the extractions from NCBI), as opposed to template-based annotations, e.g. with the MITOS algorithm (Bernt et al., 2013). This limits the precision of gene delimitation for detecting the start and stop codons and the resulting nucleotide alignment. In conclusion, the current MMG data neither help nor disturb the tree, but it is likely that after thorough taxonomic identification and improved sequence editing we will see a positive impact on the tree generated from the incongruous NCBI data, ameliorating the effect of highly uneven gene coverage.

### Current state of the Coleoptera tree

Data mining of the NCBI database provided a much greater number of taxa than the MMG datasets. However, among the >40,000 species represented at the time of data extraction (October 2015), a large proportion were represented only by a single short fragment, while just under 16,000 and 12,000 terminals, respectively, were represented by a minimum of two or three markers. Tree searches were challenging given the large number of taxa and high proportion of missing data. Yet, it was possible to apply the RAxML software (Bernt et al., 2013; Stamatakis, 2014), which is more reliable than methods designed for even larger trees, such as FastTree (Price et al., 2010), even if the algorithm is not very robust to compositional biases and high rate variation that are prevalent in mitogenome data (Pons et al., 2010; Sheffield et al., 2009; Timmermans et al., 2015b). The RAxML software permits data partitioning which improves the likelihood scores for trees from mitogenomes, whereby a split into six separate partitions for the protein coding genes (the three codon positions for forward and reverse strands) is known to provide nearly the same benefit as the separation of each gene and codon position (Lanfear et al., 2014; Timmermans et al., 2015b). Individual searches took approximately one week with our moderate local computing equipment or the CIPRES server. We observed that searches for the min3loci datasets were much faster than the min2loci, indicating that even adding a quarter more taxa and increasing the proportion of missing data put much greater demand on the searches. A search that also included species represented by only a single locus, mostly corresponding to the barcode *cox1* gene, added some 15,000 taxa to the matrix (for a total of >30,000 taxa), which was beyond the power of existing likelihood algorithms (Izquierdo-Carrasco et al., 2011), and our attempts to perform RAxML analyses on this dataset failed. Our largest tree using the min2loci dataset was obtained by using the tree from the min3loci dataset as starting tree. Thus, the greater information content (fewer missing data) of the min3loci data could be incorporated into the larger dataset without applying topological or backbone constraints based on the smaller set. This distinguishes our approach from similar efforts for species-level phylogenetics in other groups of insects, e.g. in Trichoptera, based on the barcode marker alone, which constrained several taxa at family and genus level to be monophyletic due to the limited power of these short sequences (Zhou et al., 2016). Finally, we combined the mitogenomes with the most widely used nuclear genes, in particular the 18S rRNA gene known to resolve basal relationships in the Coleoptera (Bocak et al., 2014). The end result is a generally defensible tree of an unprecedented magnitude, which now constitutes the first global assessment of relationships in all major lineages of Coleoptera (Suppl. File S3).

When evaluating the tree topology, it is tempting to focus on the basal relationships of the major subgroups, but the sequencing and sampling strategies used here are probably not the most appropriate for resolving this aspect. We consider the greatest value of the current tree of 16,000 species as a resource for the study of sublineages, e.g. within families or subfamilies. Given the high tRI, the trees are generally congruent with the existing taxonomy, which is remarkable given the rapid alignment and tree searches at the scale of many thousand taxa. This finding also implies that particularly at the highest taxonomic levels there are almost no errors in the database itself, i.e. the great majority of GenBank entries is accurate, at least at the level of families and above. We corrected a few known or suspected errors prior to the final tree searches (Suppl. File S1). The tree integrates data from independent studies that were focused on internal relationships within families or subfamilies. The relationships within these groups (discussed in detail for the families of Adephaga in the Suppl. File S1) were generally in agreement with the original studies, but the finer relationships inevitably differ because these studies may have used additional loci and more complex models.

Inconsistencies with the existing analyses were also due to non-overlapping gene sequences, in particular where either the nuclear rRNA or mitochondrial markers were available without any shared loci in these terminals or close relatives. For example, in the five representatives of the myxophagan genus *Sphaerius* (CM1S4Sp7) complete mitogenomes were available for two of the terminals, while two others were sequenced only for 18S rRNA and another one for *cox1a* and *ef1a.* Although some entries may correspond to close relatives, either gene marker placed the sequences in very different clades (the 18S-only terminals were even placed outside of Myxophaga, producing the only case that renders any of the suborders as non-monophyletic) (Suppl. File S1). To some extent the effect of non-overlapping genes can also be observed for the two subclades of Trachypachidae (the North American genus *Trachypachus* and the South American *Systolosoma*), which were placed in distant positions at the base of Carabini and Broscini, respectively. *Systolosoma* had only been sequenced for 18S rRNA and none of the mitochondrial markers, thus being attracted to other species sequenced exclusively for this gene, such as Broscini. In contrast, in *Trachypachus* the 18S rRNA was complemented by a complete mitogenome (Suppl. File S1), which allows placement relative to other mitochondrial sequences.

Another source of inconsistency is the naming of taxa. The myxophagan *Sphaerius* again illustrates a case where inconsistent naming at the species level creates extra terminals. None of the five terminals were identified to species in NCBI but instead were labeled with different alphanumeric identifiers. These specimens may correspond to the same species or even to the same specimen (e.g. submitted by the same research group as part of different datasets), and because our bioinformatics pipeline treats these as separate species entries, inflating the total number of terminals, and also placing these terminals in distant positions due to the affinities of the various gene markers.

Further problems resulting from labelling of GenBank entries are due to the casual application of taxonomic ranks and the resulting discrepancies with the Linnaean taxonomy on which the tRI is based. This was particularly evident at the genus and subgenus ranks that were not applied consistently in the NCBI classification (and the original literature on which this scheme is based). For example, the Cicindelidae, and specifically the subtribe Cicindelina, has been subdivided into subgroups by Rivalier (1953) that have been interpreted to imply a hierarchical scheme of relationships, but this is inconsistently followed by modern taxonomists. Whereas some studies have recognized two main genera named *Cicindela* and *Cylindera*, besides several other ‘genera’, the extent and internal subgeneric subdivision of these two lineages remain largely unstudied. Inevitably the mixed application of certain names at both subgeneric and generic levels in the classification inflates the tRI measure.

Finally, the heavy reliance on mitogenomes caused problems of tree inference, due to well understood problems with rate and compositional heterogeneity affecting this marker. For example, mitochondrial genomes generally recover distorted basal relationships in Adephaga, with the highly divergent aquatic families Noteridae + Meruidae separated from all others at the basal node, which renders the remaining aquatic families (Hydradephaga) paraphyletic. This well-known phenomenon (Timmermans et al., 2015a) also is reflected in the current analysis. Only additional taxon sampling or expanded sequencing of the nuclear genome will resolve these problematic basal nodes conclusively.

## Conclusions

This is the largest compilation of phylogenetic information for the Coleoptera to date. Various portions of the tree are now open to further investigation and data curation, to remove remaining errors in naming, classification, identification and sequencing. We noted in particular the errors from spelling of names and from different classificatory level to which names were attributed, which led to splitting of taxa into multiple entries in this database. Careful curation of the sequence database and the associated taxonomy database therefore is required, through requests for corrections at the NCBI database and the removal of erroneous sequence data or misassigned species. The tree enables us to identify the taxa which are placed in conflict with their identification. While some terminals may be misidentified or mislabeled, an unexpected position in certain cases can point to alternative phylogenetic relationships. The tree also offers a powerful framework for phylogenetic placement of mitochondrial reads or contigs from additional insect metagenomes or any type of environmental sequencing, facilitating biodiversity discovery.

Yet, we still have a long way to go to achieve a stable species-level tree of the Coleoptera. We show here how MMG may be a new avenue for rapidly increasing the species representation and also adding tree support. Vice versa, linking these MMG data to the existing phylogenetic database also contributes to the characterization of samples from poorly known ecosystems, such as tropical rainforests. As these data are added to the growing database, the phylogenetic placement of ecological mixtures and the generation of a more secure tree topology go hand-in-hand. Due to the unconstrained searches and inclusion of very large numbers of taxa, our approach opens the possibility for discovery of new lineages, without prior notions of taxon choice and relationships. This includes the ability to reveal new deep lineages that have hitherto been overlooked by classical taxonomy (Andujar et al., 2016; Bocak et al., 2016). This kind of sequencing effort of samples from diverse ecological and biogeographical provenances will iteratively enrich the reference tree, for an increasingly complete picture of the Tree-of-Life. Considering that the Coleoptera is the largest metazoan order on Earth, building such a reference would be a major milestone and can lead the way for similar studies across the Tree-of-Life.

## Acknowledgement

This work was funded by the Biodiversity Initiative of the NHM and NERC grant NE/M021955. PA received funding through a Newton Fellowship of the Royal Society. Thanks are due to Alex Aitken, Stephen Russell, Kevin Hopkins and Peter Foster (all NHM) for technical assistance.

## Authors contribution

BL, APV conceived the study; ACP, JM, MJTNT, CA, PA, KEM, JL, EM, AH, CGR, CB, RN, CPDTG, TB, LB collected the specimen and/or obtained the molecular data; BL, ACP and APV analysed the data; BL, APV wrote the manuscript, and all the authors revised the manuscript.

